# Date milk enriched with vitamin D: nutrient content and acceptability as a food additive for preschoolers 48-59 months

**DOI:** 10.1101/2023.05.15.540750

**Authors:** Nurnashriana Jufri, Sri Anna Marliyati, Faisal Anwar, Ikeu Ekayanti

## Abstract

Products that combine liquid milk and date flesh are still minimal. Milk is an excellent source of protein and dates are a food ingredient that is rich in vitamins and minerals that are suitable for growth as well as phytochemical components which function to enhance the sensory properties of dates so that they can be used as a flavor enhancer in various dairy products such as cookies, yogurt, ice cream, and cakes. The purpose of this study was to create a vitamin D-enriched date milk product as an alternative nutritional supplement for preschool children aged 48-59 months. The study used a completely randomized design with three percentage addition of date flesh treatments, namely F1 (10%), F2 (15%), and F3 (20%). The results showed that the formulas significantly differed between the water, protein, and carbohydrate contents (P < 0.05). In contrast, the energy, ash, fat, zinc, iron, and calcium content were not statistically significant (P > 0.05). Kruskal-Wallis analysis product acceptance data showed only color and aroma attributes that were quite different between formulas (P < 0.05). In contrast, taste, thickness, mouthfeel, aftertaste, and overall characteristics were not statistically different (P > 0.05). Formula F2 (15%) was selected based on the hedonic rating and ranking tests. Formula F2 (15%) can be accepted quite well, with the percentage that consumes drinks without residue as much as 60% and those who can finish at least ½ portion of drinks as much as 93.3%.

## Introduction

Children’s rapid growth and development during preschool require adequate and balanced nutritional intake to optimize physical growth and emotional intelligence (1). At preschool age, children begin to learn to interact with the broader environment, causing changes in eating patterns. Children are becoming active consumers who can choose what foods they like or dislike(2). This condition causes children to have the bad behavior of picky eating, namely consuming foods they only like without considering the nutritional content of these foods, which has a detrimental impact on nutritional status and health (3).

Nutritional problems in preschoolers are not only undernourished but also short. Based on Basic Health Research (Riskesdas), stunting in Indonesia is 30.8%, with a prevalence in the preschool group with concise and straightforward criteria of 7.7% and 19.2%, respectively (4). One effort to deal with nutritional problems that can be made is to develop additional food formulas that can help fulfill the nutritional needs of preschoolers (5). Foodstuffs that can be developed into additional food formulas, namely milk and dates, become a date milk drink beneficial for linear growth.

Milk has long been known to be beneficial for growth due to its protein, vitamins (A, B1, B2, E), calcium, phosphorus, magnesium, zinc, and selenium minerals. Milk is also an excellent source of protein consisting of whey protein (20% of protein) milk and casein (80% of milk protein) (6). Whey is a water-soluble protein consisting of globular proteins such as beta-lactoglobulin, alpha-lactalbumin, and the amino acids leucine, isoleucine, valine, and lysine. At the same time, casein is a water-insoluble protein containing histidine, methionine, phenylalanine, and proline which are higher (7). Casein affects growth through the process of bone mineralization by increasing the absorption of calcium from the intestine (8). Glycosylate lactoferrin, iron, and calcium-binding protein (Ca2+) in whey are physiological regulators of bone growth (9).

Since the vitamin D content of fresh milk typically ranges from 0.13-1.0 g/L and 0.03-1.86 g/kg in organic fresh milk, milk is considered a poor source of vitamin D (10). In many countries, vitamin D fortification of milk is successful in increasing vitamin D intake (11). Furthermore, vitamin D fortification is required to maximize milk’s growth benefits. Vitamin D encourages bone health at all stages of life by controlling bone remodeling and regulating phospho-calcium metabolism (11). Rickets is a bone formation and mineralization disease in infants and children due to the lack of calcium and vitamin D during bone development. Clinical symptoms of rickets include bone deformations affecting the entire skeleton, incredibly long bones, and metaphyseal growth cartilage; severe rickets disorders can result in dwarfism (12).

Dates are a fruit that contains various nutrients, including carbohydrates (85% simple sugars, namely glucose, fructose, and sucrose), protein (methionine and cysteine), complex B vitamins, such as thiamine niacin, riboflavin, pantothenic acid, pyridoxine, and folate as well as vitamin K, but has a shallow vitamin C content (13). In addition, dates are also a source of the minerals calcium, iron, cobalt, fluorine, copper, magnesium, potassium, sodium, manganese, phosphorus, sodium, zinc, sulfur, boron, and selenium (14). Consuming 100 grams can provide 15% of the RDA for the minerals copper, selenium, magnesium, and potassium, as well as 7% of the RDA for calcium, iron, manganese, and phosphorus (15).

Dates are rich in phytochemical components, including flavonoids, sterols, phenolics, anthocyanins, carotenoids, and procyanidins. In addition to pharmacological benefits, date phytochemical constituents contribute to dates’ nutritional and sensory properties, allowing dates to be used as a flavor enhancer in a variety of dairy products, including ice cream, yogurt, cookies, and cakes (16). Generally, the ingredients included in these dairy products are syrup or date juice, not fruit flesh or date flesh (17). Products that combine date flesh with milk are still quite limited in development. The purpose of this research was to produce a drink of date milk fortified with vitamin D using liquid milk and dates from the Sukkari variety, analyze the nutritional content of the formula developed, analyze the organoleptic properties, determine the selected product, and analyze the acceptability of date milk products as an alternative food additive in preschool-aged children 48-59 months.

## Methods and Materials

### Research design

This experimental study has a single factor and a completely randomized design. The research was conducted in January-February 2022. Formulation and organoleptic tests of vitamin D-enriched date milk drinks were conducted at the Food Experiment Laboratory and Sensory Analysis Laboratory, Department of Nutrition, IPB University. The MBrio Food Laboratory analyzed chemical properties, and the nutrients calcium, iron, vitamin D, and zinc were conducted at the Saraswati Indo Genetech Laboratory in Bogor. The acceptability test was conducted at the Posyandu in the Abeli and Nambo Health Centers, Kendari city working area.

### Materials and tools

The main ingredients used to produce vitamin D-enriched date milk are UHT liquid milk in tetra pack packaging and Carboxy Methyl Cellulose (CMC) emulsifier obtained from supermarkets, Dates of the Sukkari variety obtained from distributors, and Dry Vitamin D3 100 CWS/AM fortification produced by DSM Nutritional Products Ltd. Switzerland. Another ingredient is distilled water to dissolve the vitamin D fortification before mixing it into the date milk.

The tools used to manufacture products are blenders, stoves, double boiler pans, filter cloths, 200 ml plastic bottles, digital kitchen thermometers, digital food scales, measuring cups, and basins. The equipment for dissolving the fortifier is a beaker glass, measuring flask, glass funnel, stainless spatula, measuring pipette, filler pipette, stirring rod, and analytical balance. The equipment for the organoleptic test included 50 ml plastic cups, small plastic spoons, serving trays, pens, labels, and organoleptic forms. Tools for analyzing nutrient content (carbohydrates, protein, fat, ash content, moisture content, vitamin D, iron, zinc, and calcium are analytical balances, empty cups, ovens, furnaces, desiccators, Kjeldahl flasks, pipettes, flasks, digestion apparatus, distillation apparatus, Erlenmeyer flask, burette, petri dish, lead paper, soxhlet extraction apparatus, fat flask, filter paper, Buchner funnel, porcelain dish, electric cooker, HPLC spectrophotometer, stirrer, vacuum evaporator. The equipment used for the acceptance test in preschool children includes 200 ml plastic bottles, napkins, tissue, and pens.

### Making Date Milk With Added Vitamin D

The manufacture of date milk drinks is based on research by Raiesi Ardali et al (18). modified by adding 10%, 15%, and 20% of dates flesh into liquid milk. The stage of making date milk starts with preparing ingredients, mixing, adding vitamin D3 fortification, pasteurization, and cooling before being put into a bottle. The initial step in preparing the material is that the dates are sorted by selecting the fruit with good quality and then washed with running water. After that, the flesh and seeds are separated. In the mixing process, all the ingredients, namely liquid milk, fleshy dates, and CMC, are then blended for ± 2 minutes so that they are smooth and all the ingredients are mixed. After that, it is filtered with a filter cloth and sterilized beforehand to separate the date dregs (19). The addition of vitamin D fortification refers to the research by Upreti et al (20). The process of adding vitamin D fortification begins with preparing a fortifying solution. A total of 0.12-gram dry vitamin D3 powder (100 CWS/AM, DSM Nutritional Products Ltd. Switzerland) was diluted into 10 ml of distilled water, and 2 ml of the fortifying solution was added to date milk to get 600 IU/200 mL per serving of vitamin D3 in date milk drink. The pasteurization process is carried out using the High-Temperature Short-Time (HTST) method, a heating process with a temperature of 75°C for a minimum of 15 seconds (21). Pasteurization is done simply using a double boil: a large pot filled with water with a height of 7.5 to 10 cm and a smaller jar filled with date milk. The bottoms of the two pots do not touch each other to minimize the possibility of scorched date milk. Date milk is cooked over medium heat while stirring so that the date milk is homogeneous and not scorched. To measure the temperature, a sterile kitchen thermometer is used. Boiled date milk is then cooled in a basin filled with cold water until it reaches 10 C (22). The cooled date milk product is put into 200 ml bottles which have been sterilized beforehand and then stored at 4 C (23). The milk formula Vitamin D enriched dates can be seen in Table 1.

**Table 1:**
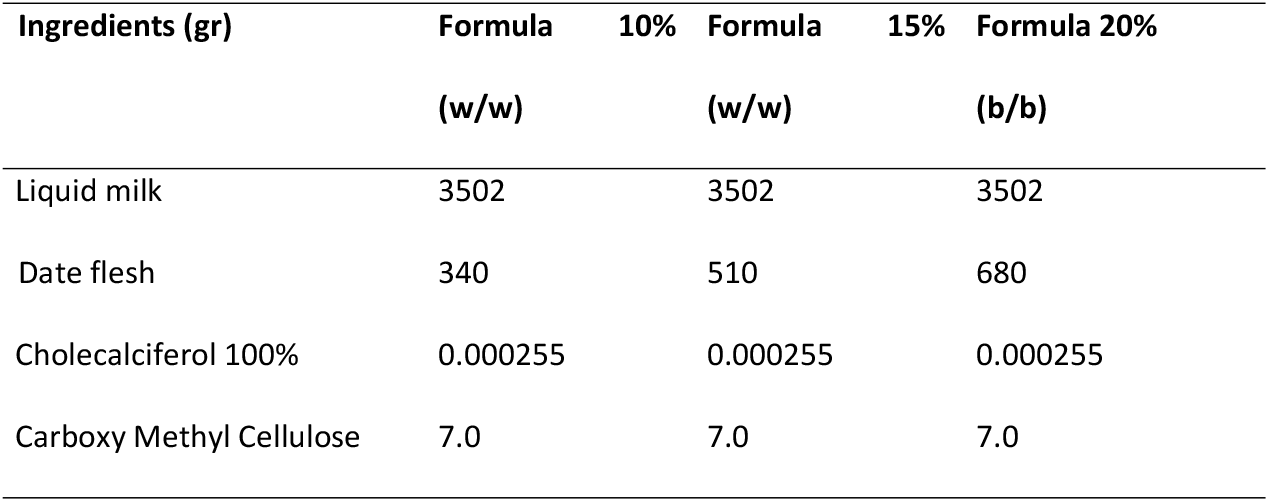
Formulation of date milk drinks fortified with vitamin D.

### Nutritional Content Analysis

Analysis of nutrient content consisted of analysis of water content and ash content using the Gravimetric method (SNI.01-2891-1992, points 5.1 and 6.1), fat content using the Soxhlet Hydrolysis method, protein content using the Kjeltech method (SNI.01-2891-1992, point 7.1), carbohydrates used the By Difference method, zinc (Zn), iron (Fe) and calcium (Ca) levels used the Inductively Coupled Plasma-Optical Emission Spectrometer (ICP OES) method. In contrast, vitamin D levels were used in the High-performance Liquid Chromatography (HPLC) method.

### Organoleptic Test and Determination of Selected Formulas

Organoleptic testing was carried out on date milk drink products enriched with vitamin D, which consisted of a hedonic rating test and a ranking test. The organoleptic test involved 30 semi-trained panelists who had received material regarding organoleptic tests or had taken organoleptic tests before and were accustomed to consuming dairy products. The hedonic rating test determines the panelists’ responses about the likes or dislikes of a product being tested through the preference level. Panelists were asked to rate their level of preference for the product on a scale of 1–7, namely 1) really disliked it; 2) do not like; 3) somewhat dislike; 4) neutral; 5) rather like; 6) likes; 7) really like the assessment attributes including color, aroma, viscosity, flavor, mouthfeel, aftertaste and overall, of a product. The ranking test was carried out to determine the panelist’s most preferred formula of the three formulas presented by rating from a scale of one to three. Panelists were asked to sort the samples by giving 1 for the most preferred product and 3 points for the least preferred product. The ranking test is carried out based on the overall attribute. The ranking values are then transformed into the scores in the Fisher and Yates Tables so that an analysis of variance can be carried out. Rank 1 scores based on the Fisher and Yates Tables have a value of 0.85, rank 2 has a value of 0, and rank 3 has a value of -0.85. The selected formula of date milk drink fortified with vitamin D was determined by choosing the best product based on organoleptic tests.

### Acceptance Test of Selected Products in Preschool Children

Acceptance test of selected formula of date milk drink fortified with vitamin D was conducted on preschool children aged 48-59 months. The acceptability test involved 30 preschool children aged 48-59 months as untrained panelists performed at several Posyandu in the Abeli and Nambo Health Centers, Kendari city, in February 2022. The acceptability test uses the Comstock method; food waste is measured by visually assessing the number of food leftovers, and the scale is based on the weighing results. In this method, the measurement scale uses the following criteria: 1) exhausted (0%); 2) Leftover ¼ portion (25%); 3) Remaining ½ portion (50%); 4) Leftover ¾ portion (75%), 5) Whole (100%).(24) The measurement result data is presented as a percentage based on a predetermined measurement scale; if the percentage of panelists consuming the product is ≥ 50%, the product is categorized as accepted by consumers. The Health Research Ethics Commission of the Institute for Research and Community Service, Halu Oleo University, approved the acceptance test for vitamin D-enriched date milk drink products with No: 1693a./UN29.20.1.2/PG/2021. Parents of children who participated in the acceptability test for vitamin D-enriched date milk products were asked to complete and sign an informed consent form to participate in this study.

### Data analysis

The IBM SPSS Statistical Analysis 24 software was used to analyze the data collected in this study. The Kruskall-Wallis test was used to analyze data on the nutritional content of each formula and organoleptic data for both the hedonic rating test and the ranking test because the data distribution was not expected. The results of the data analysis were statistically significant at the 5% level (P<0.05), and the Mann-Whitney U test was used for additional testing.

## Results and Discussion

### Nutrient content

The nutritional content of the vitamin D-enriched date milk beverage product was determined through proximate analysis, which included water content, ash content, fat, protein, and carbohydrates. At the same time, the range of zinc (Zn), iron (Fe), and calcium was analyzed using the ICP OES spectrometer method. Table 2 shows the results of the nutritional content analysis of date milk drinks enriched with vitamin D.

**Table 2:**
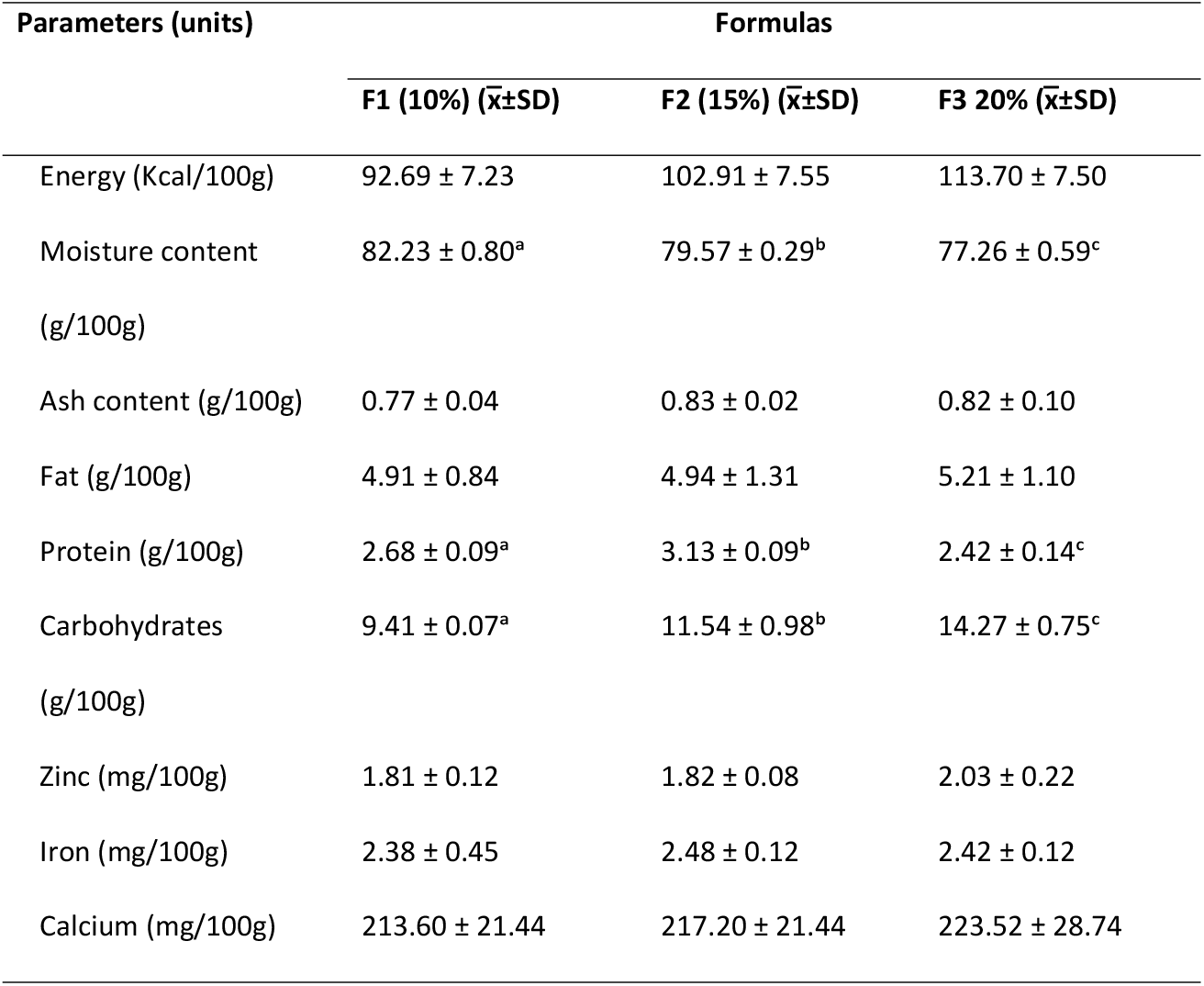
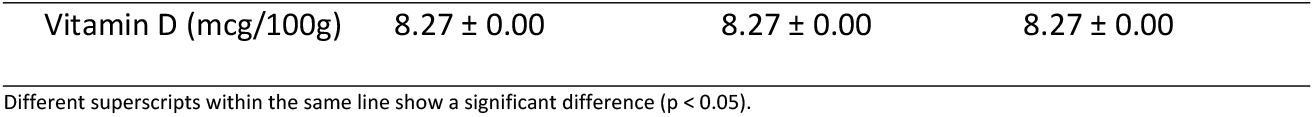
The nutritional content of the formula for date milk drinks enriched with vitamin D.

| Parameters (units) | Formulas |  |  |
| --- | --- | --- | --- |
| | F1 (10%) ($\bar{x} \pm SD$) | F2 (15%) ($\bar{x} \pm SD$) | F3 20% ($\bar{x} \pm SD$) |
| Energy (Kcal/100g) | 92.69 ± 7.23 | 102.91 ± 7.55 | 113.70 ± 7.50 |
| Moisture content<br>(g/100g) | 82.23 ± 0.80<sup>a</sup> | 79.57 ± 0.29<sup>b</sup> | 77.26 ± 0.59<sup>c</sup> |
| Ash content (g/100g) | 0.77 ± 0.04 | 0.83 ± 0.02 | 0.82 ± 0.10 |
| Fat (g/100g) | 4.91 ± 0.84 | 4.94 ± 1.31 | 5.21 ± 1.10 |
| Protein (g/100g) | 2.68 ± 0.09<sup>a</sup> | 3.13 ± 0.09<sup>b</sup> | 2.42 ± 0.14<sup>c</sup> |
| Carbohydrates<br>(g/100g) | 9.41 ± 0.07<sup>a</sup> | 11.54 ± 0.98<sup>b</sup> | 14.27 ± 0.75<sup>c</sup> |
| Zinc (mg/100g) | 1.81 ± 0.12 | 1.82 ± 0.08 | 2.03 ± 0.22 |
| Iron (mg/100g) | 2.38 ± 0.45 | 2.48 ± 0.12 | 2.42 ± 0.12 |
| Calcium (mg/100g) | 213.60 ± 21.44 | 217.20 ± 21.44 | 223.52 ± 28.74 |
| Vitamin D (mcg/100g) | 8.27 ± 0.00 | 8.27 ± 0.00 | 8.27 ± 0.00 |

Different superscripts within the same line show a significant difference ( $p < 0.05$ ).

Based on the proximate analysis performed on the three formulas, it was found that the formula with the addition of date flesh by 20% had the highest energy level, namely 113.70 Kcal/100g, and the lowest in the procedure with the addition of date flesh by 10%, namely 92.69 Kcal/100g. The energy content of the formulas did not differ significantly (P > 0.05) according to the Kruskall-Wallis test.

The energy content value in date milk drinks enriched with vitamin D was obtained by calculating fat, protein, and carbohydrate content conversion (25). Inadequate levels of energy intake are a risk factor for various nutritional problems such as malnutrition and stunting. Setiawan et al. (2018) discovered a significant relationship between caloric intake and the prevalence rate of stunting in children aged 24 to 59 months in Padang (26). Date, a fruit of the Sukkari variety, used as a raw material for the formulation of date milk drink enriched with vitamin D, has a reasonably high energy content, namely 342 kcal per 100 grams of date flesh (27). The percentage of added date flesh affects the energy content of date milk drinks enriched with vitamin D. The more significant the percentage of added date flesh, the higher the energy content.

Table 2 shows that the 10% formula has the most water content, namely 82.23 g/100g, while the lowest water content is found in the 20% formula, which is 77.26 g/100g. The difference in the additional percentage of 10%, 15%, and 20% date flesh resulted in a difference in the water content of the three formulas. Based on Kruskal-Walli’s test, it was statistically significant (P < 0.05). The statistical analysis was then continued with the Mann-Whitney follow-up test, which revealed statistically significant differences, respectively, F1 and F2, F1 and F3, and F2 and F3. Every percentage increase in the addition of date flesh produces less water content. The formula developed in this study shows that products with high energy will have low water content. The higher a product’s density and energy content, the lower its water content (28). A low water content in a product will help reduce food spoilage and prolong the shelf life of a product (25).

The formula with the highest ash content is the 15% formula of 0.83 g/100g, and the lowest is the 10% formula of 0.77 g/100g. The statistical analysis results showed no significant difference (P > 0.05) in the ash content respectively formulas using the Kruskal-Walli test. The ash content of a food ingredient can indicate its mineral content and the cleanliness and purity of the food product (29). In this study, the outcomes of the ash content analysis are produced by each fluctuating formula. In line with previous research, a decrease in the addition of date juice and an increase in the acquisition of soybean juice resulted in a fluctuating ash content in the product being developed (30). Analysis of the ash content of a food product can be used to measure food safety because it can describe the level of metal contamination in a food ingredient (31). There is a relationship between water and mineral content in food. The high-water content will reduce the mineral content because the mineral content will dissolve into the water (30). This was proven when the 10% formula had the highest water content and the lowest ash content in this study.

Analysis of the fat content in the three formulas showed that the 20% formula had the highest fat content, 5.21 g/100g, and the lowest in the 10% formula, 4.94%. The fat content of the procedures did not differ significantly (P > 0.05) according to the Kruskal Wallis test. The percentage of date flesh addition influences each formula’s resulting fat content. The addition of 20% date flesh has the highest fat content. High-fat content is expected for this product because date milk drinks enriched with vitamin D will be used as an alternative food supplement for preschoolers aged 48-59 months to support their growth. The recommended fat adequacy rate for children aged 48-59 months is 50 grams daily (32). Aside from being a supply of energy, fat is also an essential nutrient for growth because fat-soluble vitamins A, D, E, and K require fat in their metabolism (33). Ernawati et al (34) discovered a positive association between fat intake and children’s nutritional status (PB/U) (P 0.05, r = 0.046), with 11.1% of children aged six months to 12 years having concise nutritional status and 18.4% having short dietary level.

The 15% formula had the most effective average protein content, as much as 3.13 g/100g, while the lowest was in the 20% formula, as much as 2.42 g/100g. The Kruskal Wallis test revealed a statistically significant difference in protein content between procedures for vitamin D-enriched date milk drinks. The Mann-Whitney tests showed statistically significant differences, respectively, F1 and F2, F1 and F3, as well as F2 and F3. The water in a food ingredient influences the measured protein content value. The less water a food ingredient contains, the increased the approximate total protein value (35). In this study, the protein content of the three formulas had fluctuating values; the F1 and F2 formulas experienced an increase in protein content as the water content decreased. Still, in the F3 formula, the protein content decreased as the water content decreased. The decrease in protein content measured in the F3 formula is thought to be caused by the pasteurization temperature exceeding the set temperature of 75° C. According to Deana et al (36), a decrease in the protein content of the Nagara bean probiotic drink was caused by temperature fluctuations during the pasteurization process, namely 70-90° C for 20 minutes (37). The time and temperature during processing using heating such as cooking, sterilization, and drying should not be excessive, only one boiling point is enough because temperatures that are too high will cause denaturation.

High protein content in a food product is needed as a nutrient that has many benefits, including as an energy source, forming and helping the metabolism of body cells, and helping glucose metabolism (38). Protein plays an essential role in growth because it can be used for tissue synthesis, bone growth, and maintenance of body functions (39). Lack of protein intake in the long term can cause nutritional problems, including energy and protein deficiency, stunting, and exacerbate micronutrient deficiencies, including vitamin A and iron (40).

Carbohydrates are aldehyde derivatives consisting of monosaccharides, disaccharides, and polysaccharides. Carbohydrates are essential in determining the properties of food ingredients, such as flavors, color, and texture (41). Starch, sugar, and glycogen are carbohydrates that can be used as an energy source because their molecules contain carbon elements that cells can use directly (42). Carbohydrates in the body metabolize fat and protein, prevent the growth of ketosis, and prevent loss of minerals and breakdown of excess body protein (43).

The carbohydrate value contained in the three formulas was the highest in the 20% formula, 14.27 g/100g, and the lowest in the 10% formula, 9.41 g/100g. The carbohydrate content of the procedures differed significantly (P 0.05) according to the Kruskall-Wallis test. The advanced Mann-Whitney test analysis found significant differences between F1 and F2, F1 and F3, as well as F2 and F3. Sukkari dates’ raw material influences the carbohydrate content of each formula. The carbohydrate content of Sukkari date flesh is 78.32 g/100gr (27). In this study, the percentage of dates added to liquid milk was directly proportional to the value of the carbohydrate content measured in each formula, the greater the percentage of added dates, the higher the carbohydrate content of the formula.

The nutrients in each food have a different role, and macronutrients act as a source of energy, maintenance of body tissues, and metabolism of glucose, protein, and fat. Micronutrients, specifically vitamins and minerals, play a role in preventing illness which has the potential to cause problems with preschool children’s growth and development (44). Nutrients such as zinc, iron, iodine, phosphorus, and calcium are micronutrients essential in growth and stunting prevention. In achieving growth, zinc is needed to phosphorylate insulin-like growth factor 1 (IGF-1) receptors and deoxythymidine kinase activity, a process that changes deoxythymidine to deoxythymidine 5′-monophosphate, a precursor required for the synthesis of DNA, protein, and collagen (45).

Iron deficiency in children impairs growth and immune response by interfering with cytokine secretion and decreasing bactericidal macrophage activity and T-cell proliferation (46). To avoid these growth disorders, adequate iron intake from food is needed. Another nutrient that must be met for optimal child growth is calcium which plays an essential role in activating enzymes involved in energy metabolism, besides that, it also plays a role in bone mineralization to prevent bone deformation due to lack of calcium intake (28).

In this study, the results of the zinc content analysis revealed that the 20% formula had the highest value of 2.03 mg/100g and the lowest in the 10% formula of 1.81 mg/100g. The iron content was higher in the 15% formula, 2.48 mg/100g, and the lowest in the 10% formula, 2.38 mg/100g. The highest calcium value was found in the 20% formula of 223.52 mg/100g and the lowest in the 10% formula of 213.60 mg/100g. The Kruskall-Wallis test revealed no significant difference (P > 0.05) in the zinc, iron, and calcium content in each 10%, 15%, and 20% formula. The zinc, iron, and calcium content in each procedure is directly proportional to the percentage of added date flesh to liquid milk. The higher the rate of added date flesh, the higher the zinc, iron, and calcium content in each formula. In addition to the zinc, iron, and calcium content from liquid milk, which is the raw material for drinking date milk enriched with vitamin D, the addition of date flesh also contributes to the value of zinc, iron, and calcium contained in the product. It is because the zinc, iron, and calcium content in Sukkari dates is relatively high, namely zinc of 1.07 mg/100g, iron 6.50 mg/100g, and calcium of 186.55 mg/100g (27, 47) vitamin D content in each formula equals an average value of 8.27 mcg per 100 ml of drink to achieve a vitamin D content of 600 IU per serving size of 200 ml.

## Organoleptic Test Results and Determination of Selected Formulas

### Hedonic Rating Test

Data from the hedonic rating test, including the attributes of color, aroma, thickness, taste, mouthfeel, aftertaste, and overall, are presented in Table 3.

**Table 3:**
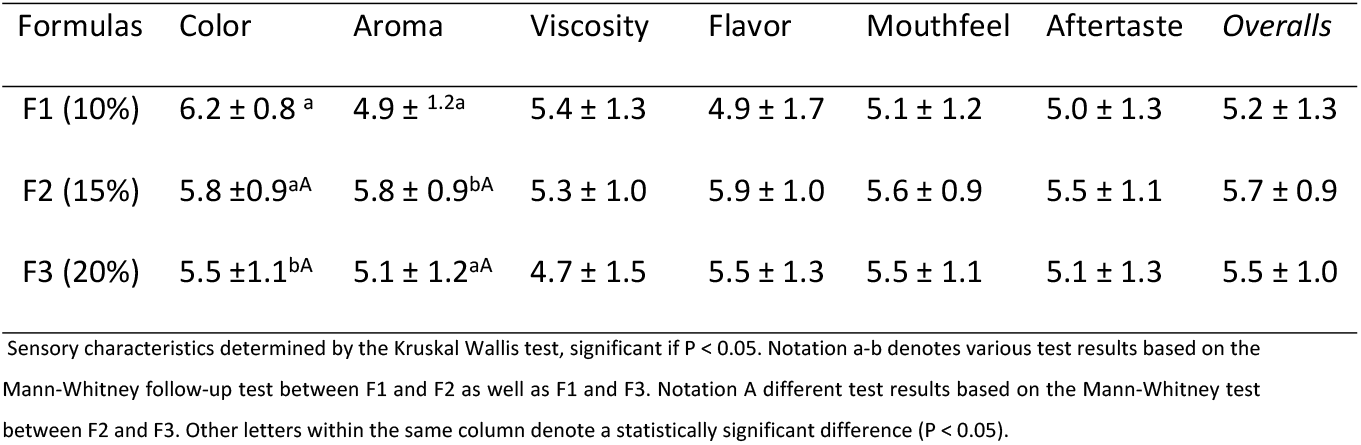
Results of the hedonic test of date milk drink enriched with vitamin D.

| Formulas | Color | Aroma | Viscosity | Flavor | Mouthfeel | Aftertaste | Overalls |
| --- | --- | --- | --- | --- | --- | --- | --- |
| F1 (10%) | 6.2 ± 0.8<sup>a</sup> | 4.9 ± 1.2<sup>a</sup> | 5.4 ± 1.3 | 4.9 ± 1.7 | 5.1 ± 1.2 | 5.0 ± 1.3 | 5.2 ± 1.3 |
| F2 (15%) | 5.8 ± 0.9<sup>aA</sup> | 5.8 ± 0.9<sup>bA</sup> | 5.3 ± 1.0 | 5.9 ± 1.0 | 5.6 ± 0.9 | 5.5 ± 1.1 | 5.7 ± 0.9 |
| F3 (20%) | 5.5 ± 1.1<sup>bA</sup> | 5.1 ± 1.2<sup>aA</sup> | 4.7 ± 1.5 | 5.5 ± 1.3 | 5.5 ± 1.1 | 5.1 ± 1.3 | 5.5 ± 1.0 |

Sensory characteristics determined by the Kruskal Wallis test, significant if $P < 0.05$ . Notation a-b denotes various test results based on the Mann-Whitney follow-up test between F1 and F2 as well as F1 and F3. Notation A different test results based on the Mann-Whitney test between F2 and F3. Other letters within the same column denote a statistically significant difference ( $P < 0.05$ ).

The first thing that attracts consumers’ attention in choosing a food product is its color appearance because the color is an attribute that consumers can directly assess through their sense of sight without having to try the product first. Based on the hedonic rating test that has been carried out, it shows that the formula with the highest average value on the color attribute is F1, namely 6.2 (likes), and the lowest is F3 with a value of 5.5 (rather likes). The Kruskal-Wallis test results revealed a difference in the percentage of added date flesh (P 0.05) on the color attribute between the formula groups. The Mann-Whitney follow-up test revealed a significant difference between F1 and F3 but no significant difference (P > 0.05) between F1 and F2 or F2 as well as F3. The results of the hedonic rating test on the color attribute can be concluded that panelists prefer F1 because it has a brighter color than F2 and F3. This color difference is influenced by the percentage of date flesh added to liquid milk, the more the portion of the number of dates added, the darker the product color will be. The dates used in the development of this formula are dates with a maturity level at the Tamar stage or also called dry dates, where the dates are very ripe, and the color of the dates turns brown to black (47). Phytochemical components found in dates, such as phenolics, anthocyanins, sterols, carotenoids, flavonoids, and procyanidins, can affect the sensory properties of dates. Anthocyanin levels are directly related to the color of the dates. In fresh dates, the anthocyanin levels are very high, while in dried dates, there is a decrease in anthocyanin levels (48).

The aroma contained in a food product is a sensory property that can be assessed using the sense of smell. The aroma attribute needs to be considered in developing a product because the aroma from one food ingredient can cause an attractive aroma or vice versa. Dates have a distinctive sweet and fresh scent (49). Based on the hedonic rating test that has been carried out, the average panelist’s perception of aroma attributes ranges from 4.9 to 5.8. The Kruskall-Wallis test and the Mann-Whitney examination revealed a significant difference (P 0.05) between F1 and F2. Still, no significant difference respectively F1 and F3, as well as between F2 and F3, so it can be concluded that there was an effect of the percentage of flesh addition dates on the aroma of date milk drink enriched with vitamin D. Dates have several volatile compounds that differ based on the type of fruit and the stage of ripening and contribute 90.7-99.6% of the total aroma profile produced. Dates contain 20 esters, 19 alcohols, 10 terpenes, 13 aldehydes, 6 ketones, 12 hydrocarbons, and one lactone as volatile compounds, which produces a characteristic citrus, floral, fruity, and herbal aroma in dates (50).

The texture is a food property that can be assessed using the eyes, skin, and mouth muscles, product acceptance indicators (49). In this study, the surface observed was the thickness of the drink being developed. The hedonic rating test that has been carried out shows that the average value of panelist perceptions on the viscosity attribute ranges from 4.7 to 5.4. The Kruskal-Wallis test results showed no difference in the percentage of added date flesh (P > 0.05) on the formula’s viscosity attribute. Nevertheless, the portion of addition of date flesh tends to affect the thickness of the procedure for date milk drinks enriched with vitamin D. The higher the percentage of added date flesh, the less preferred date milk products because the product’s viscosity level is higher. Beverage products with a liquid texture are generally preferred over thick drinks. In line with a study conducted by Violeta and Mardiana (2022), which showed that formulas with a runny consistency are selected because the surface is like mineral water which does not cause discomfort to the throat when the product is consumed (51).

Taste is a biological perception in the form of taste sensations (sweet, bitter, sour, salty, astringent, cold, and hot) produced by matter received by the sense of taste through taste receptors in the mouth and also accepted by aroma receptors in the nose. Taste can be raised through the aroma of food ingredients, allowing the tongue to feel other flavors besides bitter, salty, sour, and sweet according to the scent (52). The outcomes of the hedonic rating test revealed that the average panelist perception value on the taste attribute ranged from 4.9 to 5.9, with formula F2 receiving the highest score. The Kruskal Wallis test showed that adding date flesh did not affect the taste of the resulting drink (P > 0.05). The F2 formula tends to be more popular because the taste is judged to be just right, not too sweet, and not bland. Date fruit used as a flavor enhancer in date milk drinks enriched with vitamin D has a varied sugar content, including monosaccharides (glucose of 51.80 g/100gr and fructose of 47.50 g/100gr) followed by disaccharides (sucrose of 3.20 g/100 gr). The Sukkari variety, used as a raw material for making beverages, has a higher glucose and fructose content than other varieties, such as the Allig dates and Deglet-Nour (27).

Mouthfeel is a complex sensation caused by a food product’s physical and chemical characteristics with texture parameters such as stringy, oily, watery, gritty, and crumbly (53). Based on statistical analysis using the Kruskal Wallis test, the treatment with adding date palm flesh did not affect the mouthfeel attribute (P > 0.05). The panelists’ average value for the mouthfeel attribute ranged from 5.1 to 5.6, indicating that the panelists’ acceptance of the developed drink formula was quite good. Formula F2 is the formula that has the highest average value, which means that this formula is preferred compared to the other two procedures. The F2 formula, adding 15% date flesh, provides a suitable mouthfeel attribute, which is neither too thick nor too runny. The process of filtering date flesh during processing is carried out twice, namely after mixing liquid milk and date flesh using a blender and at the stage before filling the date milk drink into packaged bottles. This process is intended to minimize the gritty and stringy sensation when the glass is consumed.

The taste sensation on the tongue after food or drink is swallowed or vomited is known as the aftertaste and is an essential factor in product quality and acceptance by consumers (54). In the aftertaste attribute, the mean value of the formula ranges from 5.0 to 5, indicating that between F1, F2, and F3, the range of panelist perception values is not too far away. However, the F2 formula has a greater panelist preference than the other two. Kruskal-Walli’s analysis revealed that adding date flesh to the treatment did not affect the product’s aftertaste attributes (P > 0.05). The use of dates as a natural sweetener in the development of vitamin D-enriched date milk drink formulas results in no excessive sensation felt by the sense of taste after the drink is swallowed.

The overall attribute is general acceptability, assessed based on observations of various attributes: color, aroma, thickness, taste, mouthfeel, and aftertaste (28). The results of the hedonic rating test on the overall attribute show an average value ranging from 5.2 to 5.7, the formula with the highest average value is F2. Based on Kruskall-Wallis test conducted showed that the difference in the percentage of adding date flesh did not affect the overall attribute (P> 0.05). It can be concluded that the three formulas are at the same level.

### Ranking Test

The data in Table 4 is the result of the ranking test based on the order of preference of the panelists. The developed formula shows that F2 has the highest score, 0.25, and the lowest in F1, -0.28. The Kruskall-Wallis test, followed by Mann-Whitney revealed a significant difference between F1 and F2 (P 0.05), but no significant difference between F1 and F3 as well as F2 and F3 (P > 0.05).

**Table 4:**
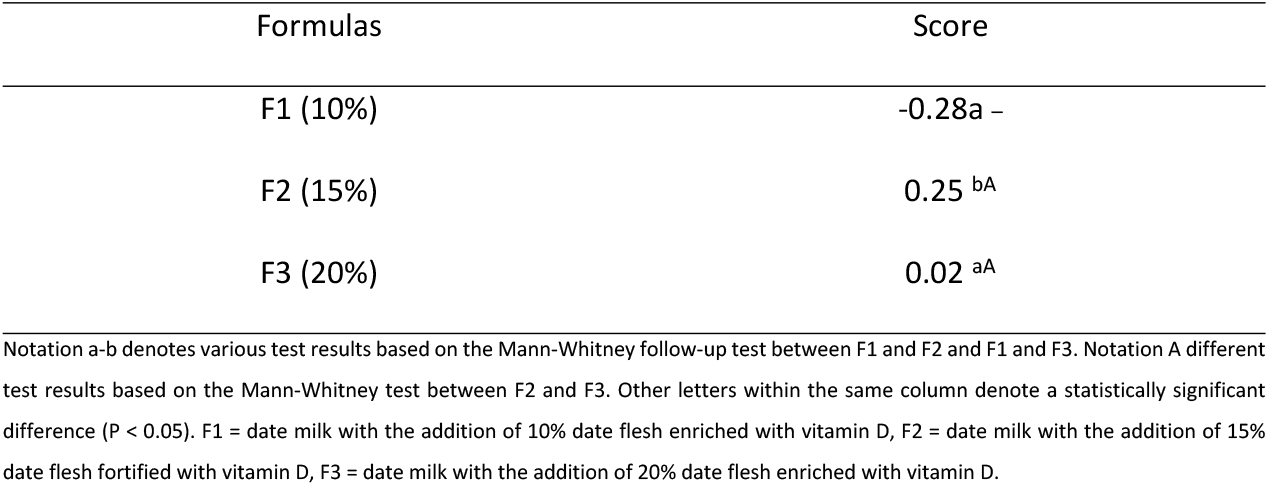
Ranking test results of date milk drinks fortified with vitamin D.

The basis for assessing the best product is looking at the highest average value of the overall attribute in the hedonic rating test. In contrast, the ranking test is done by accumulating the ranking values for each formula, and the product with the highest score value has opted for the best and preferred outcome. Based on the results of the hedonic rating and ranking tests, it can be concluded that the formula chosen based on the panelists’ preference level was F2, namely the procedure for drinking date milk with the addition of 15% flesh of dates enriched with vitamin D.

### Acceptance Test of Selected Products in Preschool Children

The acceptability test for selected products was conducted on preschool children aged 48-59 months using the Comstock form. Acceptance criteria are measured by looking at the remaining drinks given. Table 5 showed the distribution of product acceptance in preschool children aged 48-59 months, revealing that there were 18 (60%) children who finished (0%) drinking date milk fortified with vitamin D, 6 (20%) children who left ¼ portion (25%), 4 (13.3%) children left ½ portion (50%), 2 (6.7%) children left ¾ portion (75%). No child left a whole drink (100%).

**Table 5:** Distribution of Acceptance of Selected Products in Preschool Children.

| Leftover Drink F2 | n | Percentage (%) |
| --- | --- | --- |
| Used up (0%) | 18 | 60 |
| Remaining $\frac{1}{4}$ portion (25%) | 6 | 20 |
| Remaining $\frac{1}{2}$ portion (50%) | 4 | 13.3 |
| Remaining $\frac{3}{4}$ portion (75%) | 2 | 6.7 |
| Whole (100%). | 0 | 0 |
| Total | 30 | 100 |

The Comstock method is the most widely used method for measuring food waste because the Comstock visual scale is based on weighing results so that the results of assessing food waste are not too different from food weighing. Nisak et al (55) discovered that the Comstock method was more efficient than the food weighing method in evaluating food waste.

The indicator of acceptance of a food product is when the percentage of consumers who reject a product is less than 50%. The acceptance test results in this study demonstrated that there were 60% of preschoolers finished the drink without residue, 20% concluded ¾ portion, and 13.3% spent ½ amount so that a total of 93.3% of children could consume at least ½ dose of vitamin-enriched date milk drink D. The sweet taste is the reason the preschooler likes the date milk drink given. The same logic was discovered in a study performed by Aini et al (56), where 63% of school-age children liked it, and 33% enjoyed a snack bar with bee pollen products with added chocolate as a sweetener and flavor enhancer, so children selected the product.

## Conclusion

The addition of 10%, 15%, and 20% date flesh had a substantial impact (P < 0.05) on the water, protein, and carbohydrate content of the drink but had no significant effect on the ash, fat, energy, zinc, iron, and calcium content of the glass (P > 0.05). Formula F2 with the addition of date flesh of 15% is the selected formula in the development of date milk drink enriched with vitamin D. The results of the nutritional content analysis of the chosen procedure include energy, moisture content, ash content, fat, protein, carbohydrates, zinc, iron, calcium, and vitamin D respectively are 102.91 kcal/100g, 79.57 g/100g, 0.83 g/100g, 4.94 g/100g, 3.13 g/100g, 11.54 g/100gr, 1.82 mg/100gr, 2.48 g/100gr, 217.20 mg/100gr and 8.27 mcg/100gr. The acceptability of the vitamin D enriched date milk drink given to preschoolers aged 48-59 months is quite good because there are 60% of preschoolers finish the drink without residue, 20% finish ¾ portion, and 13.3% spend ½ pore, so a total of 93.3% of children who can consume at least ½ portion and which leaves ¾ portion only 6.7%.

## Acknowledgments

The author would like to thank all of the panelists who took part in the organoleptic test for the development of vitamin D enriched date milk products, as well as the posyandu cadres in the Nambo and Abeli health center work areas who assisted carry out the acceptability test of selected vitamin D enriched date milk products in preschool children.

## References

1. Hardianti R, Dieny FF, Wijayanti HS. Picky eating dan status gizi pada anak prasekolah (Picky eating and nutritional status in preschool children). Jurnal Gizi Indonesia (The Indonesian Journal of Nutrition). 2018;6(2):123–30.

2. Afrinis N, Indrawati I, Raudah R. Hubungan Pengetahuan Ibu, Pola Makan dan Penyakit Infeksi Anak dengan Status Gizi Anak Prasekolah (Correlation between Mother’s Knowledge, Diet, and Children’s Infectious Diseases with Nutritional Status of Preschool Children). Aulad: Journal on Early Childhood. 2021;4(3):144–50.

3. Chao HC. Association of picky eating with growth, nutritional status, development, physical activity, and health in preschool children. Front Pediatr. 2018;6(February):1–9.

4. Kemenkes RI. Laporan Nasional Hasil Riset Kesehatan Dasar (Riskesdas) Indonesia tahun 2018. Riset Kesehatan Dasar 2018. 2018. p. 166.

5. Irwan Z, Salim A, Adam A, Irwan Z, Salim A, Adam A. Giving cookies of Moringa leaf flour and Moringa seed flour towards weight and nutritional status of children in the Tampa Padang public health center. Jurnal AcTion: Aceh Nutrition Journal. 2020;2020(5):45–54.

6. Yackobovitch-Gavan M, Phillip M, Gat-Yablonski G. How milk and its proteins affect growth, bone health, and weight. Horm Res Paediatr. 2017;88(1):63–9.

7. Haug A, Høstmark AT, Harstad OM. Bovine milk in human nutrition - A review. Lipids Health Dis. 2007;6(25):1–16.

8. Holt C, Carver JA, Ecroyd H, Thorn DC. Invited review: Caseins and the casein micelle: Their biological functions, structures, and behavior in foods1. J Dairy Sci. 2013;96(10):6127–46.

9. Grenov B, Michaelsen KF. Growth Components of Cow’s Milk: Emphasis on Effects in Undernourished Children. Food Nutr Bull. 2018;39(2_suppl): S45–53.

10. Mandrioli M, Boselli E, Fiori F, Teresa Rodriguez-Estrada M. Vitamin D3 in high-quality cow milk: An Italian case study. Foods. 2020;9(5):1–9.

11. Marangoni F, Pellegrino L, Verduci E, Ghiselli A, Bernabei R, Calvani R, et al. Cow’s Milk Consumption and Health: A Health Professional’s Guide. J Am Coll Nutr. 2019;38(3):197–208.

12. Allgrove J, Shaw NJ. A Practical Approach to Vitamin D Deficiency and Rickets. In: Endocrine Development. 2015. p. 119–33.

13. Al-Farsi MA, Lee CY. Nutritional and functional properties of dates: A review. Crit Rev Food Sci Nutr. 2008;48(10):877–87.

14. Al-Farsi M, Alasalvar C, Al-Abid M, Al-Shoaily K, Al-Amry M, Al-Rawahy F. Compositional and functional characteristics of dates, syrups, and their by-products. Food Chem. 2007;104(3):943–7.

15. Abdeen E sayed MM. Enhancement of Functional Properties of Dairy Products by Date Fruits. Egypt J Food. 2018; 46:197–206.

16. Hussain MI, Farooq M, Syed QA. Nutritional and biological characteristics of the date palm fruit (Phoenix dactylifera L.) – A review. Food Biosci. 2020; 34:100509.

17. Keshtkaran M, Mohammadifar MA, Asadi GH, Nejad RA, Balaghi S. Effect of gum tragacanth on rheological and physical properties of a flavored milk drink made with date syrup. J Dairy Sci. 2013;96(8):4794–803.

18. Raiesi Ardali F, Rahimi E, Tahery S, Shariati MA. Production of a New Drink by Using Date Syrup and Milk. Journal of Food Biosciences and Technology. 2014;4(2):67–72.

19. J. K. Negara, M. Arifin, E. Taufik, T. Suryati. Penambahan Sari Kurma sebagai Substrat Antibakteri pada Minuman Whey Fermentasi. Jurnal Ilmu Produksi dan Teknologi Hasil Peternakan. 2021;9(1):36–41.

20. Upreti P, Mistry V V., Warthesen JJ. Estimation and fortification of vitamin D3 in pasteurized process cheese. J Dairy Sci. 2002;85(12):3173–81.

21. Triwidyastuti Y, Nizar M, Jusak J. Temperature Control in Milk Pasteurization Process with Temperature Control Milk Pasteurization Utilizing the Proportional-Integral-Derivative (Pid) and Fuzzy Sugeno Method. Jurnal Teknologi Informasi dan Ilmu Komputer (JTIIK). 2019;6(4):355–62.

22. Yuni S, Madanijah S, Setiawan B, Marliyati SA. Pengembangan Produk Yang Berpotensi Sebagai Minuman Fungsional Untuk Penderita Prahipertensi (Product Development with Potential as a Functional Drink for Prehypertension Sufferers). Jurnal Gizi dan Pangan. 2016;11(2):135–42.

23. Nzekoue FK, Alesi A, Vittori S, Sagratini G, Caprioli G. Development of functional whey cheese enriched in vitamin D3: nutritional composition, fortification, analysis, and stability study during cheese processing and storage. Int J Food Sci Nutr. 2021;72(6):746–56.

24. Manjilala, Ekariskawati, Agustian I. Daya terima bolu cukke substitusi tepung kulit pisang dan tepung tempe pada balita gizi kurang (Acceptability of cukke cake as a substitute for banana skin flour and tempeh flour in undernourished toddlers). Media Gizi Pangan. 2019;26(1):71–7.

25. Hastuti AR, Afifah DN. Analysis of Antioxidant Activity, Analysis of Nutritional Content, Organoleptic Tests of Sesame Seed Snack Bars and Pumpkin Flour as Alternative Snacks with High Antioxidants. Journal of Nutrition College. 2019;8(4):219–30.

26. Setiawan E, Machmud R, Masrul M. Faktor-Faktor yang Berhubungan dengan Kejadian Stunting pada Anak Usia 24-59 Bulan di Wilayah Kerja Puskesmas Andalas Kecamatan Padang Timur Kota Padang Tahun 2018 (Factors Associated with Stunting Incidents in Children Aged 24-59 Months in the Working Area of Andalas Health Center, East Padang District, Padang City in 2018). Jurnal Kesehatan Andalas. 2018;7(2):275.

27. Siddeeg A, Zeng XA, Ammar AF, Han Z. Sugar profile, volatile compounds, composition and antioxidant activity of Sukkari date palm fruit. J Food Sci Technol. 2019;56(2):754–62.

28. Mentari AD, Setiawan B, Palupi E. Pengembangan RUTF (Ready to Use Therapeutic Food) Berbahan Serealia dan Kedelai bagi Balita Malnutrisi Akut Berat (Development of RUTF (Ready to Use Therapeutic Food) Made from Cereals and Soybeans for Severe Acute Malnutrition Toddlers). Media Gizi Indonesia. 2022;17(1):11–20.

29. Palupi E, Rahmatika M. Peningkatan Nilai Gizi Pada Susu Tempe Kedelai Hitam (Glycine soja sieb) (Improvement of Nutritional Value of Tempeh Milk Made from Black Soybean (Glycine soja sieb)). Jurnal Gizi Dietetik. 2022;1(1):42–9.

30. Sabariman M, Wahyuningtias ES, Azni IN. Formulasi jus kurma dan sari kedelai dalam pembuatan jus kurma soya (Formulation of date juice and soy juice in the manufacture of soy date juice). Jurnal Teknologi Pangan dan Kesehatan. 2022; 4(1): 55–66.

31. Kristiandi K, Rozana R, Junardi J, Maryam A. Analysis of Moisture, Ash, Fiber, and Fat Content in Siam Orange Syrup Drink (Citrus nobilis var. microcarpa). Jurnal Keteknikan Pertanian Tropis dan Biosistem. 2021; 9(2):165–71.

32. Kemenkes RI. Peraturan Menteri Kesehatan Republik Indonesia Nomor 28 Tahun 2019 Tentang Angka Kecukupan Gizi yang Dianjurkan Untuk Masyarakat Indonesia (Regulation of the Minister of Health of the Republic of Indonesia Number 28 of 2019 concerning Recommended Nutrition Adequacy Rates for Indonesian People). Jakarta; 2019 p. 12–4.

33. Siregar FA, Makmur T. Metabolisme Lipid Dalam Tubuh (Lipid Metabolism in the Body). Jurnal Inovasi Kesehatan Masyarakat. 2020; 1(2):60–5.

34. Ernawati F, Yuriestia Arifin A, Prihatini M. Hubungan Asupan Lemak dengan Status Gizi Anak Usia 6 Bulan-12 Tahun di Indonesia (Relationship of Fat Intake with Nutritional Status of Children Aged 6 Months-12 Years in Indonesia). Penelitian Gizi dan Makanan. 2019;42(1):41–7.

35. Normilawati, Fadlilaturrahmah, Hadi S, Normaidah. Penetapan Kadar Air dan Kadar Protein pada Biskuit Yang Beredar Di Pasar Banjarbaru (Determination of Moisture Content and Protein Content in Biscuits Circulating in the Banjarbaru Market). Jurnal Ilmu Farmasi. 2019;10(2):51–5.

36. Yusuf IE, Swamilaksita PD, Ronitawati P, Fadhilla R. Pengembangan Tepung Sukun dan Tepung Kacang Tunggak dalam Pembuatan Kue Mangkok (Development of Breadfruit Flour and Cowpea Flour in Making Cupcakes). Jurnal Pangan dan Gizi. 2022 Apr 14;12(1):71–82.

37. Deana O, Agustina L, Musthafa Al Hakim H. Probiotik Kacang Nagara (Vigna unguiculata ssp. Cylindrica) [Effect of Type and Stabilizer Concentration on the Manufacture of Probiotik Drinks from Nagara Beans (Vigna unguiculata ssp. Cylindrica)]. Pro Food (Jurnal Ilmu dan Teknologi Pangan) [Internet]. 2019 Nov;5(2):496–506. Available from: http://www.profood.unram.ac.id/index.php/profood

38. Sukini T. Efektivitas Konsumsi Nugget Tempe Kedelai Terhadap Kenaikan Berat Badan Balita Gizi Kurang (The Effectiveness of Consuming Soybean Tempeh Nuggets on Weight Gain in Undernourished Toddlers). Jurnal Kebidanan. 2017;6(12):63–72.

39. Uauy R, Kurpad A, Tano-Debrah K, Otoo GE, Aaron GA, Toride Y, et al. Protein and Amino Acids in Infant and Young Child Nutrition. J Nutr Sci Vitaminol. 2015; 61:192–4.

40. Agustina M, Rimbawan R, Setiawan B, Herminiati A. Pengaruh Pemberian Diet Rendah Protein dan Restriksi Pakan pada Pertumbuhan dan Protein Serum Tikus Lepas Sapih (Effect of Low Protein Diet and Feed Restriction on Growth and Serum Protein in Weaning Mice). Nutri-sains: Jurnal Gizi, Pangan dan Aplikasinya. 2019;5(1):1–14.

41. Fitri AS, Fitriana YAN. Analisis Senyawa Kimia pada Karbohidrat (Analysis of Chemical Compounds in Carbohydrates). Sainteks. 2020;17(1):45–52.

42. Maulana Setiawan, Wiratama I, Sulaeman A. Peranan Karbohidrat Dalam Perspektif Alqur’an (The Role of Carbohydrates in the Perspective of the Qur’an). Educatoria: Jurnal Ilmiah Ilmu Pendidikan. 2022;2(4):249–57.

43. Kole H, Tuapattinaya P, Watuguly T. Analysis of Carbohydrate and Fat Levels in Tempe Made from Seagrass Seeds (Enhalus acoroides). BIOPENDIX: Jurnal Biologi, Pendidikan dan Terapan. 2020;6(2):91–6.

44. Mayar F, Astuti Y. Peran Gizi Terhadap Pertumbuhan dan Perkembangan Anak Usia Dini (The Role of Nutrition on Early Childhood Growth and Development). Jurnal Pendidikan Tambusai. 2021;5(3):9695–704.

45. Brion LP, Heyne R, Lair CS. Role of zinc in neonatal growth and brain growth: review and scoping review. Pediatr Res. 2021;89(7):1627–40.

46. Domellöf M, Braegger C, Campoy C, Colomb V, Decsi T, Fewtrell M, et al. Iron requirements of infants and toddlers. J Pediatr Gastroenterol Nutr. 2014;58(1):119–29.

47. Hussain MI, Farooq M, Syed QA. Nutritional and biological characteristics of the date palm fruit (Phoenix dactylifera L.) – A review. Food Biosci. 2020;34(December 2019):1–12.

48. Al-Farsi M, Alasalvar C, Morris A, Baron M, Shahidi F. Comparison of antioxidant activity, anthocyanins, carotenoids, and phenolics of three native fresh and sun-dried date (Phoenix dactylifera L.) varieties grown in Oman. J Agric Food Chem. 2005;53(19):7592–9.

49. Lababan FMJ, Rahmawati YD. Uji Daya Terima dan Nilai Gizi Bolu Kukus yang Disubstitusi Kurma (Phoenix Dactylifer) sebagai Altenatif Jajanan Pencegahan Anemia (Test of Acceptability and Nutritional Value of Steamed Spongebob Substituted with Dates (Phoenix Dactylifer) as an Alternative Snack to Prevent Anemia). Jurnal Ilmiah Gizi Kesehatan 2022;3(02):82–8.

50. Siddiq M, Aleid SM, Kader AA. Date Fruit Composition and Nutrition. In: Siddiq M, Aleid SM, Kader AA, editors. Dates: Postharvest Science, Processing Technology and Health Benefits. First Edit. USA: John Wiley & Sons Ltd; 2014. p. 261–83.

51. Violeta D, Mardiana M. Antioxidant Levels and Preference Test for Moringa Leaf and Date Fruit Combination Drinks to Improve Athlete’s Performance. Journal of Nutrition College. 2022;11(4):328–36.

52. Tarwendah IP. Studi Komparasi Atribut Sensori dan Kesadaran Merek Produk Pangan. Jurnal Pangan dan Agroindustri (Comparative Study of Sensory Attributes and Brand Awareness of Food Products. Journal of Food and Agroindustry). 2017;5(2):66–73.

53. Guinard JX, Mazzucchelli R. The sensory perception of texture and mouthfeel. Trends Food Sci Technol. 1996;7(7):213–9.

54. Schlossareck C, Ross CF. Consumer sensory evaluation of aftertaste intensity and liking of spicy paneer cheese. Int J Food Sci Technol. 2020;55(7):2710–8.

55. Nisak NK, Ronitawati P, Citra K. PDAT and Comstock Methods Are More Efficient Than Food Weighing in Assessing Patient Food Leftovers. Nutrire Diaita. 2019;11(1):18.

56. Aini Q, Sulaeman A, Sinaga T. Development of Bee Pollen Snack Bars for School-Age Children. Jurnal Teknologi dan Industri Pangan. 2020;31(1):50–9.

